# Gastric Neurons in the Nucleus Tractus Solitarius are Selective to the Orientation of Gastric Electrical Stimulation

**DOI:** 10.1101/2021.07.09.451818

**Authors:** Jiayue Cao, Xiaokai Wang, Terry L. Powley, Zhongming Liu

## Abstract

Gastric electrical stimulation (GES) is a bioelectric intervention for gastroparesis, obesity, and other functional gastrointestinal disorders. In a potential mechanism of action, GES activates the nerve endings of vagal afferent neurons and induces the vago-vagal reflex through the nucleus tractus solitarius (NTS) in the brainstem. However, it is unclear where and how to stimulate in order to optimize the vagal afferent responses. To address this question with electrophysiology in rats, we applied mild electrical currents to two serosal targets on the distal forestomach with dense distributions of vagal intramuscular arrays that innervated the circular and longitudinal smooth muscle layers. During stimulation, we recorded single and multi-unit responses from gastric neurons in NTS and evaluated how the recorded responses depended on the stimulus orientation and amplitude. We found that NTS responses were highly selective to the stimulus orientation for a range of stimulus amplitudes. The strongest responses were observed when the applied current flowed in the same direction as the intramuscular arrays in parallel with the underlying smooth muscle fibers. Our results suggest that gastric neurons in NTS may encode the orientation-specific activity of gastric smooth muscles relayed by vagal afferent neurons. This finding suggests that the orientation of GES is critical to effective engagement of vagal afferents and should be considered in light of the structural phenotypes of vagal terminals in the stomach.

## Introduction

The vagus is the primary neural pathway that connects the brain and the stomach (Browning & Travagli, 2011; Levinthal & Strick, 2020; Powley et al., 2019). It enables rapid stomach-brain interactions (Andermann & Lowell, 2017; Carabotti et al., 2015; Furness and Stebbing, 2018; Holtmann & Talley, 2014; Mayer, 2011; Williams et al., 2016) and reflexive control of the gastrointestinal tract (Babic & Browning, 2014; Travagli & Anselmi, 2016). Neural signaling via the vagus is bi-directional. The afferents ascend sensory signals to the nucleus tractus solitarius (NTS) (Rogers et al., 1995; Powley & Phillips, 2002). The efferents descend motor commands from the dorsal motor nucleus of the vagus (DMV) (Travagli et al., 2006). Despite progress in structural characterization of vagal projections to the stomach (Brookes et al., 2013; Powley et al., 2019), very little is known about the specific gastric information transmitted to and encoded by the brain. In particular, it is unclear how NTS, as the first nucleus for visceral sensation, represents and integrates gastric information relayed through vagal afferents.

Both vagal afferents and efferents innervate the stomach with extensive and specialized terminals in the stomach wall (Brookes et al., 2013; Furness et al., 2020; Powley & Phillips, 2002). The afferent terminals exhibit distinctive morphologies and chemical codes sensitized to various gastric signals (Berthoud et al., 1997; Berthoud & Powley, 1992; Powley et al., 2019). In particular, the intramuscular array (IMA) runs in parallel with muscle fibers and serves as mechanoreceptors to report on the stretch (or length) of gastric smooth muscles (Berthoud & Powley, 1992; Wang & Powley, 2000; Phillips & Powley, 2000; Powley et al., 2011). It has two subtypes, namely longitudinal and circular IMAs, which target longitudinal and circular muscles, respectively. Both subtypes show regional distributions overlapping in the antrum but distinctive in the forestomach (or fundus) (Powley et al., 2016; Tan et al., 2021).

Stimulation targeted to vagal afferent terminals is one mechanism of gastric electrical stimulation (GES) (Cheng et al., 2021). Electrical current pulses with a low amplitude, high frequency, and short duration are inadequate to activate the muscle or pace the stomach, but can potentially relieve nausea and vomiting and treat motor dysfunctions for drug-resistant gastroparesis (Abell et al., 2003; Lal et al., 2015; Levinthal & Bielefeldt, 2017; McCallum et al., 2010; O’Grady et al., 2009). Despite its approved use in clinics, the efficacy of GES varies across patients and its parameterization remains empirical and suboptimal. A potential strategy to optimize GES is to engage the central nervous system via vagal afferent signaling (Gottfried□Blackmore et al., 2020; Mayer et al., 2006; McCallum et al., 2010; Payne et al., 2019; Qin et al., 2005; Tang et al., 2006). However, it is unclear where and how to apply GES in order to maximize the vagal afferent responses.

In this study, we targeted GES to longitudinal and/or circular IMAs and recorded neuronal responses from gastric nuclei in NTS. Since IMAs were orientation specific (Powley et al., 2016), we hypothesized that the GES-evoked vagal afferent responses were dependent on the stimulus orientation. To test this hypothesis in rats, we chose two serosal targets in the distal forestomach with high IMA densities and applied short current stimuli with different orientations and amplitudes. We recorded the neuronal spiking responses in NTS and evaluated the responses across various stimulus settings. By testing this hypothesis, we aimed to characterize the functional specialization of gastric neurons in NTS and to offer additional clues as to how to apply GES in order to modulate the stomach-brain neuroaxis.

## Materials and Methods

### Subjects

This study used seven Sprague–Dawley rats (male, weight: 250–400□g; Envigo RMS, Indianapolis, IN) according to a protocol approved by the Institutional Animal Care and Use Committee and the Laboratory Animal Program at Purdue University. All animals were housed in a strictly controlled environment (temperature: 21□±□1□°C; 12□h light-dark cycle with light on at 6:00 a.m. and off at 6:00 p.m.). Every animal received acute surgical implantation of patch electrodes for gastric electrical stimulation and depth electrodes for neuronal recordings in the brainstem.

### Gastric electrical stimulation

Every animal received abdominal surgery to be implanted with patch electrodes on the serosal surface of the distal forestomach. To prepare for the experiment, the animal was fasted overnight one day before the experiment. Food was removed from the cage at 6 pm, while the experiment started at 10 AM the next day. The overnight fasting ensured a nearly empty stomach during the experiment. The animal was initially anesthetized with 5% isoflurane mixed with oxygen (flow rate: 1 L/min). The dose of isoflurane was then reduced to 2% to keep a surgical plane. Then, the animal was placed in a supine position on the surgical bench. Following a toe-pinch test, an incision was made starting from the xiphoid and moving 4□cm caudally. Skin and muscle layers were retracted and separated to expose the ventral forestomach.

Patch electrodes (Microprobes, Gaithersburg, MD, USA) were sutured at two target locations, namely Target 1 and 2, on the serosal surface of the distal forestomach (figure 1(a)). Target 1 was on the greater curvature and about 4 mm away from the limiting ridge. Target 2 was on the ventral stomach, 2 mm away from the limiting ridge, and halfway in between the lesser curvature and the greater curvature. According to our previously mapped distributions of longitudinal and circular IMAs (Powley et al., 2016; Tan et al., 2021), Target 1 had a high density of longitudinal IMAs but a low density of circular IMAs, whereas Target 2 had a high density of circular IMAs but a low density of longitudinal IMAs (figure 1(b)). Target 1 and Target 2 were chosen as complementary targets selective to the “hotspots” of longitudinal and circular IMA, respectively.

**Figure 1.**
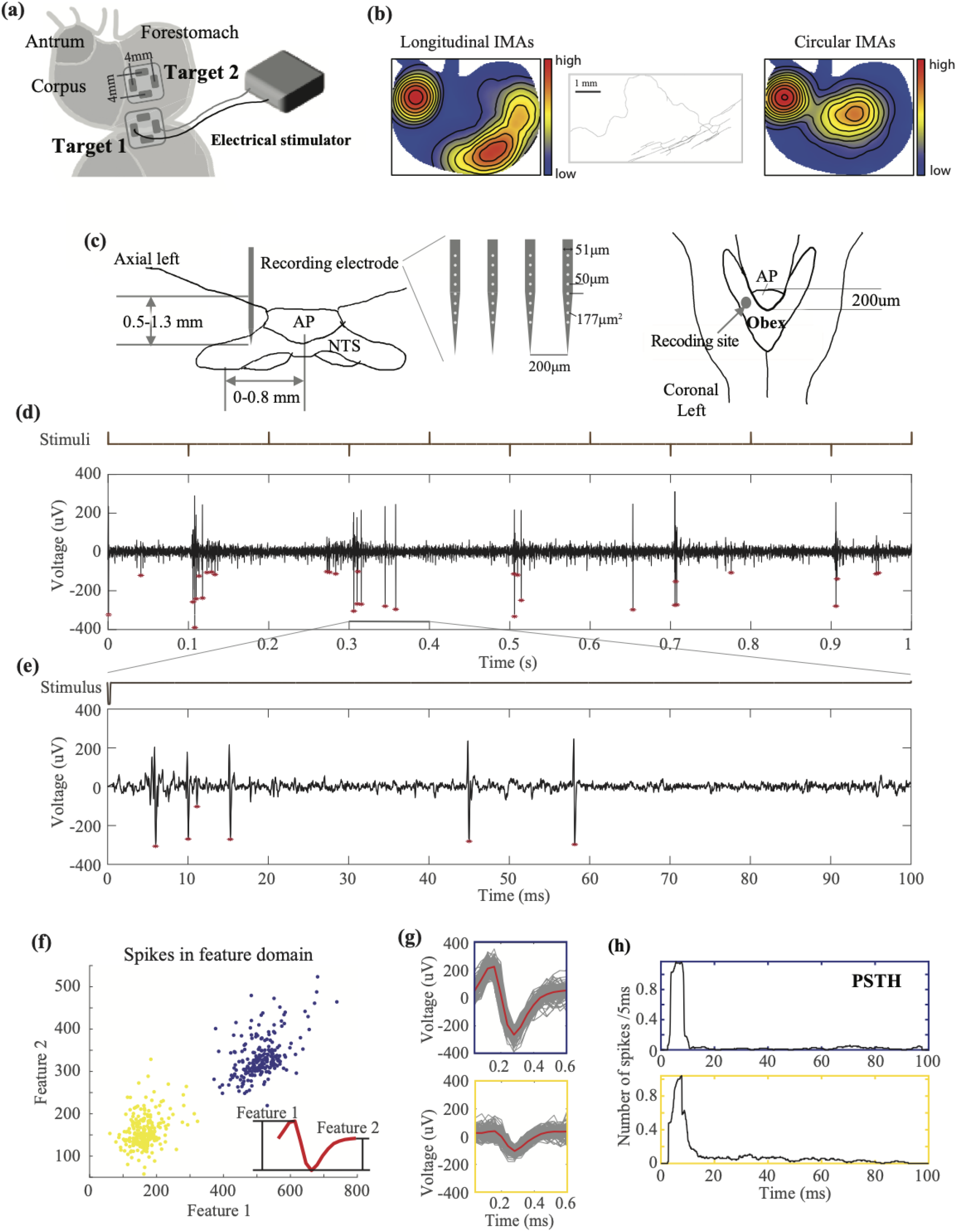
Experiment protocol and analysis pipeline. **(a)** shows the experimental setup and the location of the implanted electrodes for gastric electrical stimulation. **(b)** shows the distributions and morphology of IMAs. The distributions of longitudinal and circular IMAs are shown in the left and right respectively (Tan et al., 2021). The middle shows an example of a parent axon branching out to parallel arrays of longitudinal IMAs. **(c)** illustrates the position of the silicon probe inserted into the NTS relative to the area postrema (AP) and Obex in the axial and coronal view, respectively. **(d)** shows an example time series of MUA (bottom) during a train of alternating current pulses (top) applied to Target 1 in the longitudinal direction. Positive pulses deliver the current toward the distal stomach. Negative pulses deliver the current toward the proximal stomach. The population spikes are marked with red stars. **(e)** shows the zoom-in of the same time series within 100 ms following a single negative pulse. The stimulus onset is at time 0. **(f)** shows the feature representations of individual spikes. Each dot is a spike described in terms of two spike features, which are illustrated in the red plot in the bottom-right inset. Two clusters of spikes are highlighted in yellow and blue. Each cluster corresponds to a single neuron. **(G)** shows the individual spikes (in grey) that belong to each cluster and their average within the cluster (in red). The boundary of the top or bottom plot is color-matched to the blue or yellow cluster, respectively. **(h)** shows the post-stimulus time histogram of the firing rate (the average number of spikes per 5 ms) for each of the two neurons sorted in **(f)** and **(g)**.

Each patch electrode consisted of either one or two bipolar pairs of contacts. Each contact was made of a Pt/Ir foil (12 mm^2^) on a thin perylene substrate (77 mm^2^, thickness = 0.18 mm) and was positioned 2 mm away from the center of the substrate. Each bipolar pair was oriented for current delivery along either the longitudinal or circular direction. We attached two pairs of electrodes on Target 1 for four animals, two pairs of electrodes on Target 2 for four animals, and one pair of electrodes on Target 1 for three animals (Table 1). We fixed the electrodes by securing the four corners of the substrate on the serosal layer. Additional sutures were used to the edges of the substrate as needed to secure the electrode contact with the serosal surface. After implantation of the electrodes, the muscle layer and the skin were closed with sutures, while the leads were kept outside of the abdomen and were connected to a current stimulator (model 2200, A-M Systems, Sequim, USA).

**Table 1.** The number and orientation of the implanted electrodes in each animal. The number of electrodes is specified with respect to the location (Target 1 vs. 2) and the direction (longitudinal vs. circular).

| Animal ID | Target 1 |  | Target 2 |  |
| --- | --- | --- | --- | --- |
|  | Longitudinal | Circular | Longitudinal | Circular |
| 1 | 2 | 2 | 0 | 0 |
| 2 | 2 | 2 | 0 | 0 |
| 3 | 2 | 2 | 0 | 0 |
| 4 | 2 | 2 | 2 | 2 |
| 5 | 2 | 0 | 2 | 2 |
| 6 | 2 | 0 | 2 | 2 |
| 7 | 2 | 0 | 2 | 2 |

For each target location, electrical current was delivered in four plausible orientations: longitudinal pointing towards the proximal (0°) or distal stomach (180°) with one pair of electrodes, or circular pointing toward the lesser (90°) or greater (270°) curvature with the other pair of electrodes (figure 2(b)). Current stimuli were delivered as a train of rectangular pulses with alternating polarity (pulse width: 0.3 ms, pulse amplitude: 0.6, 0.8, or 1 mA). Specifically, one positive monophasic pulse was first delivered, followed by one negative pulse. The interval between the two adjacent (positive or negative) pulses was 100 ms (10 Hz) for two animals and 250 ms (4 Hz) for the other five animals. This pattern was repeated, with the pulse train lasting 10 s followed by a stimulus-free rest period of >20 s. This 10s pulse train was repeated 3 to 5 times for each recording session. Different pulse trains were configured to stimulate in either the longitudinal or circular direction in an order randomized and counterbalanced across animals.

**Figure 2.**
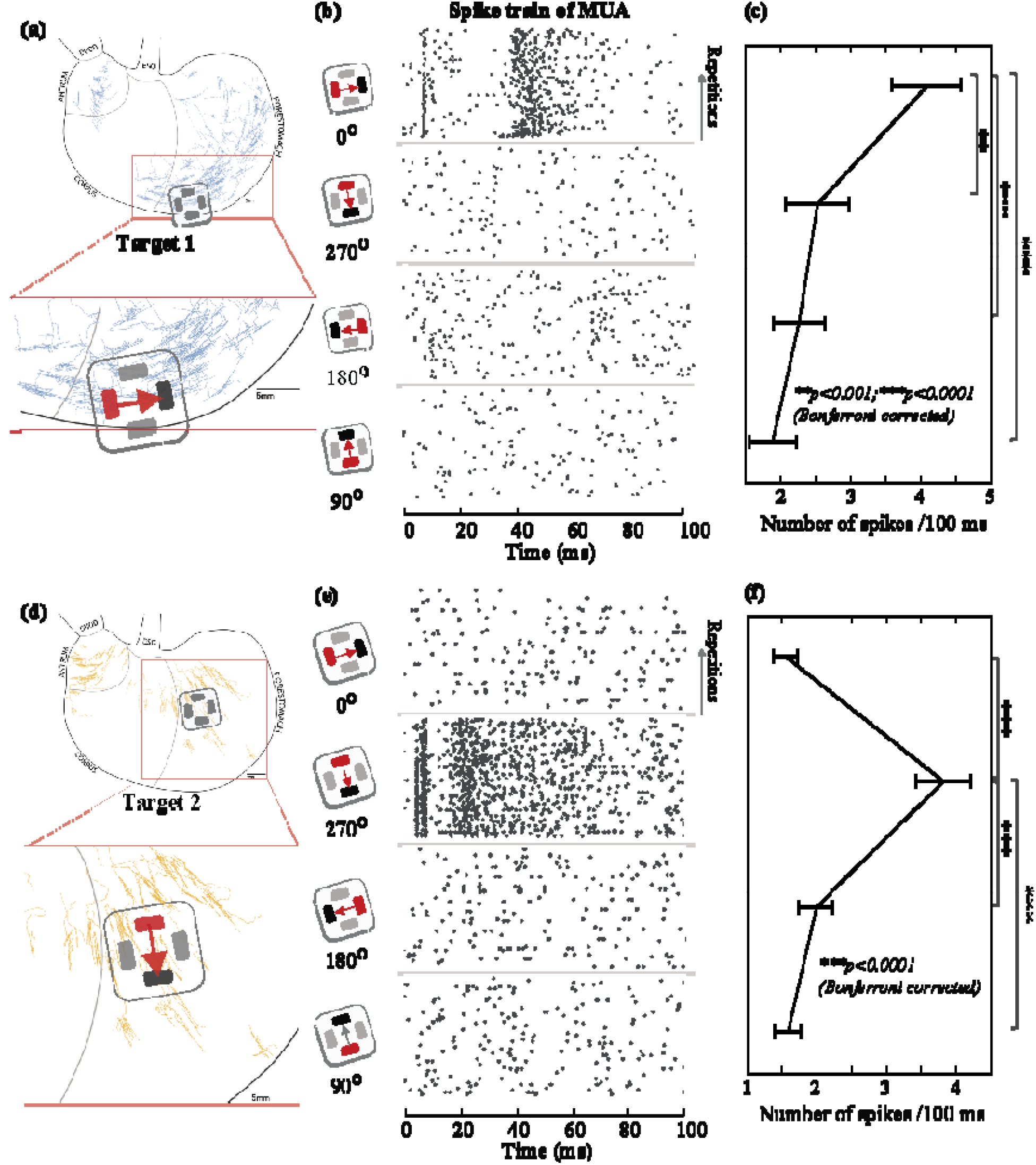
GES-evoked population spikes in NTS given different stimulus orientations. **(a)** and **(d)** show the locations of GES in comparison with the digitized samples of longitudinal and circular IMA based on prior work (Powley et al., 2016; Tan et al., 2021). **(b)** and **(e)** show the pairing of the anode (red) and the cathode (black) for delivering a current pulse in one of four different directions (0°, 270°, 180°, and 90°). In **(b)** (or (**e)**), the raster displays the timing of the population spikes observed from one channel in response to a single current pulse across repeated trials. Each black dot marks the time of a spike. Each row indicates a single trial. Time 0 is the stimulus onset. Data in **(b)** and **(e)** are from different channels. **(c)** (or **(f)**) plots the average number of spikes during the (100 ms) post-stimulus period given each orientation, after averaging across trials. The error bar indicates the standard error of the mean. ** means *p*<0.001 and *** means *p*<0.0001 for paired t-test with Bonferroni correction.

### Neurophysiological recording

To record neuronal spiking activity, we used a 32-channel silicon probe (Figure 1.C; Neuronexus, Ann Arbor, USA; product number: A4×8-5mm-50-200-177). Briefly, the silicon probe includes four shanks. The four shanks are separated by 200 μm. Each shank has eight channels, as shown in Figure 1.C. The distance between adjacent channels is 50 μm, and the area of each channel is 177 μm2. Each individual shank has a width of 51 μm and a thickness of 15 μm.

To implement the electrodes in each animal, a surgery was performed by placing the animal in the prone position on a stereotaxic frame (Stoelting Co., Wool Dale, IL, USA). To expose the skull, a midline skin incision was made from between the eyes to the neck. Two stainless steel screws were drilled into the skull above the olfactory bulb and were used as the ground and reference. A 5×5 mm^2^ cranial window was created at the bottom of the occipital bone to expose the brainstem until the obex was identifiable. Right before the recording session, a bolus of dexdomitor (12.5 μg/Kg, 0.05mg/ml; zoetis, Parsippany-Troy Hills, New Jersey, USA) was administered subcutaneously, followed by continuous subcutaneous infusion of dexdomitor at 12.5 μg/(Kg x hour) (0.05 mg/ml). During the infusion, isoflurane was reduced to 0.1-0.5% mixed in oxygen to maintain the normal physiological condition. The silicon probe was inserted at about 0-0.8mm left and 0-0.2mm rostral to the obex (figure 1(c)). The depth of insertion was determined by using a searching process to identify gastric neurons in NTS. Specifically, gastric stimulation in all possible orientations was delivered (pulse width = 0.3 ms, current = 1.0 mA, pulse frequency = 1 Hz) while the probe was being inserted into NTS. The insertion was stopped once the stimulus-evoked spiking activity was observable in one or multiple channels, regardless of the orientation of the stimuli delivered. The searching process was repeated multiple times to identify different groups of neurons, while each time the probe was pulled out and re-inserted to different depths and coordinates (from 0.5 to 1.3 mm). At each depth of insertion, neural data was recorded from all 32 channels by using a broadband recording system (Tucker Davis Technologies, Alachua, USA) with a sampling rate of 24 kHz for a total of 1 to 2 hours.

### Single and multi-unit firing rate

We applied a high-pass filter (>300 Hz) to the raw signals to extract the neuronal spiking activity. First, unsorted spikes were identified if they surpassed a threshold (between 50 and 100 □V) defined for each animal. The unsorted spikes were stored by channel as the multi-unit activity (MUA). Second, spike sorting was applied separately to each channel to extract the single-unit activity (SUA). The spike sorting extracted from each spike two features based on the peak-trough differences: the difference between the first peak and the trough (Feature 1) and the difference between the second peak and the trough (Feature 2). As illustrated in figure 1(f), the spike features were used to group every spike into distinct clusters where each cluster corresponded to a neuron. The clustering was based on the k-means algorithm (kmeans in Matlab), where the distance between spikes was their Euclidean distance in the feature space. The number of clusters was determined based on the distributions of spike representations in the feature space. The sorted spikes across all channels were stored as SUA.

Based on the SUA or MUA, we evaluated the neuronal responses to GES in the level of either single or multiple units, respectively. For each neuron, the SUA response was quantified as the number of spikes within a 100-ms post-stimulus period in each trial of GES parameterized by its location (Target 1 & 2), direction (0°, 90°, 180°, 270°), and amplitude (0.6, 0.8, 1 mA). The single-trial SUA response was averaged across repeated stimuli with the same parameters and then across animals. Similarly, the time-averaged MUA response was quantified for each channel.

For a given location and amplitude of GES, the time-averaged MUA response was compared across four stimulus orientations (0°, 90°, 180°, 270°). The orientation that resulted in the highest MUA response was considered as the preferred orientation for each channel under evaluation. Then, the MUA response was averaged across channels and animals, to obtain the group averaged NTS responses as a function of the current amplitude (0.6, 0.8, 1 mA).

In addition to the time-averaged SUA or MUA response, we further evaluated the response as a function of the post-stimulus latency. The time-resolved response was based on the post-stimulus time histogram (PSTH). Specifically, the number of spikes was calculated within a 5 ms sliding window stepping by one time point (1/24 kHz) until reaching 100 ms after the stimulus. The firing rate (i.e., the number of spikes per 5ms) was averaged across repeated trials of the same stimulus setting. Neurons (for SUA) or channels (for MUA) that showed their maximal PSTH less than 0.05 spikes per 5 ms were discarded and excluded from subsequent analyses. Further, the PSTH was averaged within four coarsely defined post-stimulus periods: 0-13 ms, 13-29 ms, 29-73 ms, and 73-100 ms.

### Statistical analysis and orientation selectivity

We used a paired t-test to evaluate the statistical significance of any response difference given GES applied in any two different orientations. The t-test was separately performed for the mean SUA or MUA response averaged over the 100-ms post-stimulus period. Bonferroni correction was used to correct for multiple comparisons (i.e., 6 pairs of different orientations under comparison). The significance level a was set as 0.05. Similarly, we also used a paired t-test to evaluate the difference between GES applied to two different locations (i.e., Target 1 and 2).

We used an orientation selectivity index (OSI) to measure how a neuron (SUA) or neuronal ensemble (MUA) was selective to GES applied in the preferred orientation vs. other orientations. Specifically, let N_PO_ be the firing rate in response to GES in the preferred orientation (e.g., 0°), and let N_i_ (i=1 to 3) be the firing rate in response to GES in the non-preferred orientations (e.g., 90°, 180 °, 270 °). The OSI was the average difference in the firing rate between the preferred orientation and every non-preferred orientation, as expressed by Equation 1. The OSI was in a range from 0 to 1, while negative values were set to 0. A higher value of OSI indicated that the neuron or channel under evaluation was more selective to the GES in the preferred orientation.

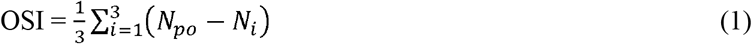

We used a permutation test to evaluate the statistical significance of the OSI. Specifically, the SUA or MUA responses were randomly shuffled into different groups defined by the stimulus orientation. The shuffled dataset disrupted any real response-stimulus association. After repeating the process 100,000 times and at each time evaluating the OSI with the shuffled dataset, we generated a null distribution of OSI, against which the actual OSI was compared for calculation of the *p-value*.

## Results

In seven rats, we recorded single and multi-unit activity from gastric neurons in NTS while applying GES to two complementary regions of interest in the distal forestomach (figure 1(a)): Target 1 with a high density of longitudinal IMA (figure 1(b) left) and Target 2 with a high density of circular IMA (figure 1(b) right), according to recent studies (Powley et al., 2016; Tan et al., 2021). We evaluated the GES-evoked neuronal responses as a function of the stimulus orientation, location, and amplitude. In particular, our emphasis was on the orientation selectivity of gastric neurons in NTS. This was motivated by the fact that the IMA, as an array of vagal afferent receptors (Berthoud and Powley, 1992), typically shows a parent arbor branching into elongated and parallel receptors directly innervating muscle fibers (Wang and Powley, 2000). Figure 1(b) middle shows a digitized example of IMA. Figure 2(a) and 2(b) further show the topographic distributions of longitudinal and circular IMA samples (Powley et al., 2016), respectively.

### Orientation dependence of NTS population responses to GES

When GES was delivered as a current pulse with a short width (0.3 ms) and a low amplitude (<1 mA), the stimuli were too mild to directly pace the stomach or induce muscle contractions (Du et al., 2009; Li & Chen, 2010; Cheng et al., 2021; Forster et al., 2001; McCallum et al., 1998). However, such GES could evoke population spikes (figure 1(d) & (e)) observable with a 32-channel silicon probe inserted into NTS (figure 1(c)). Figure 1(d) shows a typical MUA recording in NTS during a train of alternating currents delivered to Target 1 in two opposing directions (i.e., positive and negative pulses). In this example, many more spikes were observed following the negative pulses than following the positive pulses (figure 1(d)). The spikes occurred shortly after each negative pulse (figure 1(e)). By spike sorting (figure 1(f)), two neurons were identified (figure 1(g)) and their responses were observed mostly in the first 20 ms following the pulse (figure 1(h)). This observation led us to hypothesize that gastric neurons in NTS were selective to the orientation of GES.

We extended the above observation by performing quantitative comparisons across four orientations and two locations, which covered the non-overlapping “hotspots” (figure 1(b)) of longitudinal and circular IMAs shown in figure 2(a) and 2(d), respectively. For Target 1, when a 1mA current stimulus was applied in 0° (i.e., flowing in the longitudinal direction towards the proximal stomach), the stimulus induced multiple population spikes with spike timing highly consistent across repeated trials (figure 2(b)). For an example MUA recorded from one channel, one spike occurred around 10 ms; a few spikes occurred from 40 to 60 ms. When the current stimuli were applied in other directions, fewer spikes were evoked and were less consistent across trials (figure 2(b)). Beyond this example, the population response in NTS, in terms of the number of spikes per 100 ms averaged across channels and animals, was 4.09±0.50 (mean ± sem) for GES at 0°, which was significantly higher than those for other orientations (2.52±0.45 for 270°; 2.26±0.37 for 180°; 1.88±0.34 for 90°), as shown in figure 2C. In summary, 1-mA GES at Target 1 evoked reliable population spikes in NTS when the stimulus was delivered in 0° (longitudinal towards the proximal stomach).

Similarly, for Target 2, we observed much stronger and more reliable population responses when 1-mA GES was applied in 270° (circular towards the greater curvature) relative to other orientations. Given the preferred orientation, figure 2(e) shows an example of the population spikes observed from one channel. The spike timing was highly consistent across repeated trials. The population spikes were fewer and much less consistent given GES in non-preferred orientations. The mean firing rate was 3.83±0.39 per 100 ms (mean ± sem) for GES at 270°, which was significantly higher than those for other orientations (1.56±0.17 for 0°; 1.99±0.25 for 180°; 1.56±0.19 for 90°) (figure 2(f)).

### Effects of the current amplitude

We asked whether the orientation selectivity of NTS neurons observed with 1-mA GES was generalizable to GES with lower amplitudes. To address this question, we compared the neuronal population responses in NTS given GES with three different current amplitudes (0.6, 0.8, and 1 mA). For both Target 1 (figure 3(a)) and Target 2 (figure 3(d)), the time-averaged population responses in NTS were always greater when GES was applied in its preferred orientation (0° for Target 1, 270° for Target 2), despite the difference in the current amplitude used in this study (figure 3(b) & (e)). In its preferred orientation, a stimulus with a higher amplitude tended to induce stronger population responses in NTS. Stimuli in other non-preferred orientations did not show significant amplitude-dependent effects on the resulting population responses.

**Figure 3.**
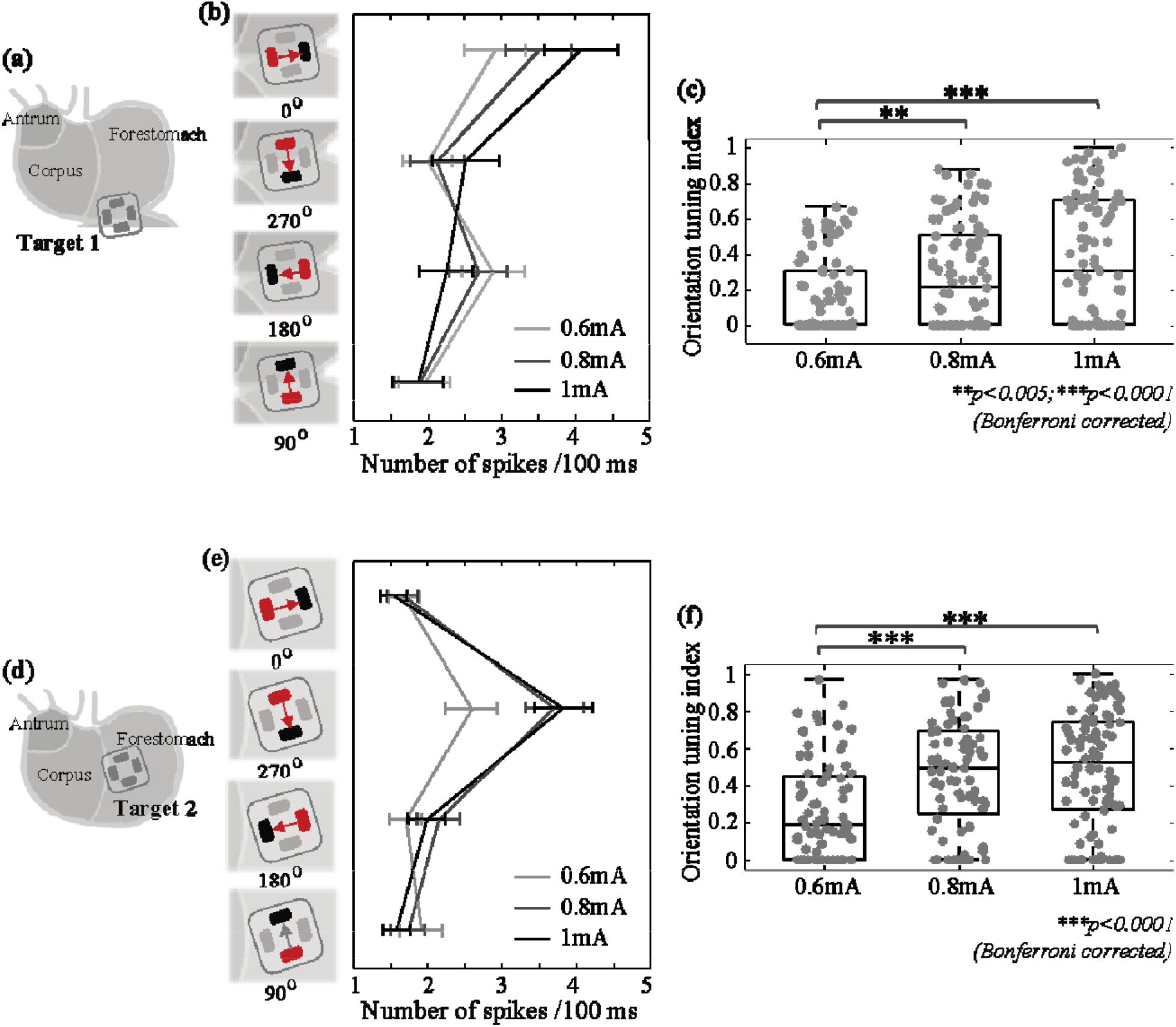
Population responses and orientation selectivity in NTS given GES with different current amplitudes. **(a)** (or **(d)**) shows the electrodes implanted to Target 1 (or 2). **(b)** (or **(e)**) shows the mean firing rate in response to GES at Target 1 (or 2) in each of the four orientations with the current amplitude at 0.6, 0.8, or 1 mA. The error bar indicates the standard error of the mean. **(c)** (or **(f)**) shows the orientation selectivity index (OSI) given different current amplitudes for Target 1 (or 2). Grey dots are individual samples. ** means p<0.005 and *** means p<0.001 (permutation test with Bonferroni correction).

We further calculated the orientation selectivity index (OSI) as the average difference in the neuronal response to GES between the preferred and non-preferred orientations described above. For both Target 1 and 2, increasing the current amplitude led to more pronounced orientation selectivity (i.e., increasingly higher OSI) (figure 3(c)). For Target 1, the OSI at 0.6 mA was significantly lower than the OSI at 0.8 mA (*p*<0.005) or 1 mA (*p*<0.001) based on a permutation test with Bonferroni correction. The observations and statistics were similar for Target 2 (figure 3(f)). See Table 2 and Table 3 for statistical results in further details. In summary, orientation selectivity was generalizable across different current amplitudes (0.6, 0.8, and 1 mA).

**Table 2.** Statistical difference in the mean population firing rate (i.e., the number of spikes per 100 ms) given GES applied to Target 1 with a preferred orientation (0°) vs. other orientations (270°, 180°, 90°). Each row corresponds to a different current amplitude. The statistics below are based on paired t-tests with Bonferroni correction for multiple comparisons.

| current | Firing rate at 0° | 0° vs 270° | 0° vs 180° | 0° vs 90° |
| --- | --- | --- | --- | --- |
| 0.6mA | 2.91±0.42 /100ms | t(91)=3.48; p=0.0023 | t(91)=0.54; p=0.59 | t(91)=3.56; p=0.0018 |
| 0.8mA | 3.51±0.45 /100ms | t(92)=4.74; p=2.3e-5 | t(92)=7.26; p=3.7e-10 | t(92)=6.32; p=2.8e-8 |
| 1mA | 4.09±0.50 /100ms | t(92)=3.82; p=7.2e-4 | t(92)=6.93; p=1.7e-9 | t(92)=7.63; p=6.2e-11 |

**Table 3.**
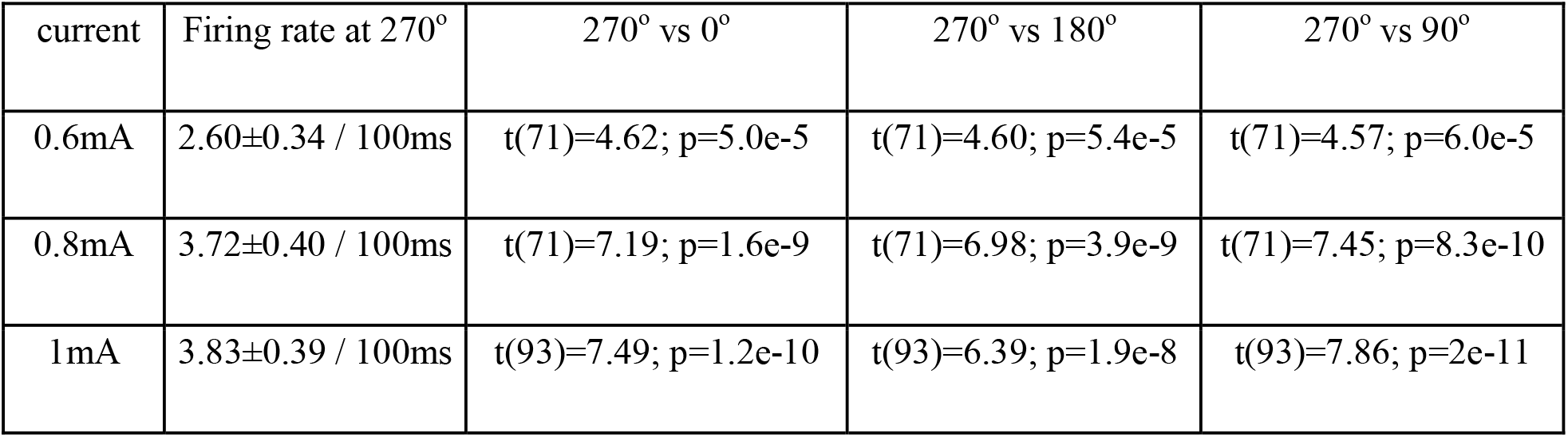
Statistical difference in the mean population firing rate (i.e., the number of spikes per 100 ms) given GES applied to Target 2 with a preferred orientation (270°) vs. other orientations (0°, 180°, 90°). Each row corresponds to a different current amplitude. The statistics below are based on paired t-tests with Bonferroni correction for multiple comparisons.

| current | Firing rate at 270° | 270° vs 0° | 270° vs 180° | 270° vs 90° |
| --- | --- | --- | --- | --- |
| 0.6mA | 2.60±0.34 / 100ms | t(71)=4.62; p=5.0e-5 | t(71)=4.60; p=5.4e-5 | t(71)=4.57; p=6.0e-5 |
| 0.8mA | 3.72±0.40 / 100ms | t(71)=7.19; p=1.6e-9 | t(71)=6.98; p=3.9e-9 | t(71)=7.45; p=8.3e-10 |
| 1mA | 3.83±0.39 / 100ms | t(93)=7.49; p=1.2e-10 | t(93)=6.39; p=1.9e-8 | t(93)=7.86; p=2e-11 |

### Orientation selectivity of single-unit responses

We further tested whether the orientation selectivity was applicable to the level of single neurons in NTS. We applied spike sorting to identify individual neurons activated by GES and analyzed their firing rates as a function of the stimulus orientation and location. In total, we identified 106 neurons activated by GES at Target 1 and 119 neurons by GES at Target 2.

Most neurons activated by GES manifested high selectivity for one stimulus orientation. In figure 4(b), an example neuron showed reliable responses only when GES at Target 1 was in 0°, whereas virtually no responses were observed for other orientations. In total, 67 out of 106 neurons all shared the same preferred orientation (i.e., 0°) and showed significantly higher firing rates given GES in 0° than in other orientations (figure 4(c)). Neurons with a preference for other orientations (i.e., 270°, 180°, 90°) were not only fewer but also less orientation selective (i.e., significantly lower OSI) (figure 4(d) & (e)). Similarly for GES at Target 2, 86 out of 119 neurons were highly selective to the same preferred orientation of 270° (figure 4(f) through (j)). In summary, gastric neurons in NTS were selective to 0° (longitudinal towards the proximal stomach) for GES at Target 1 and were selective to 270° (circular towards the greater curvature) for GES at Target 2.

**Figure 4.**
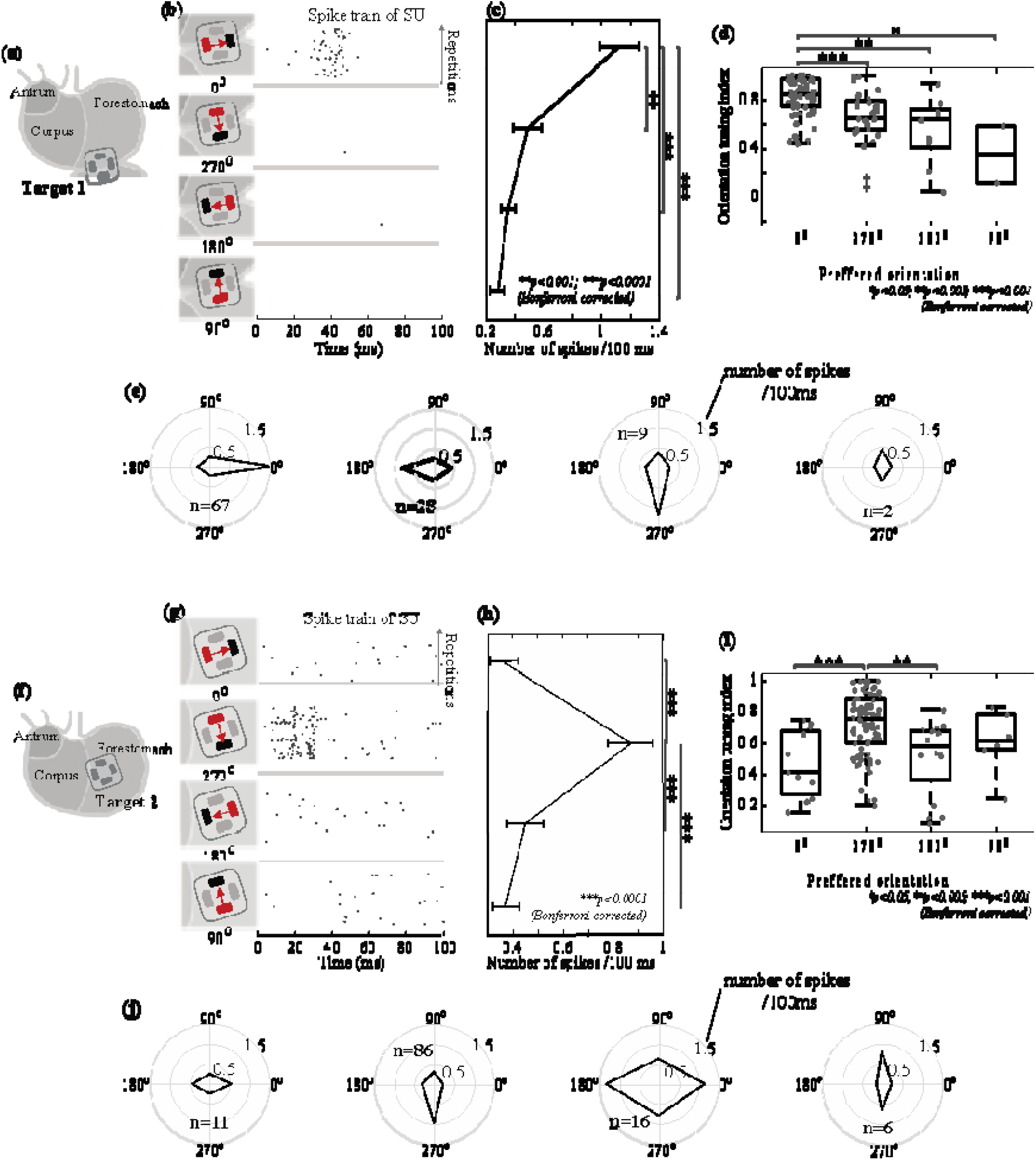
Orientation selectivity for single neurons in NTS. For 1-mA GES at Target 1 **(a)** and Target 2 (**f**), **(b)** and **(g)** show the firing of a neuron (right) in response to a current pulse delivered in four different orientations (left). The raster displays the timing of SUA across repeated trials. Each dot marks the time of a spike. Each row is one stimulus trial. Time 0 is the stimulus onset. Data shown in **(b)** and **(g)** are from different neurons. **(c)** and **(h)** plot the average firing rate in terms of the number of spikes per 100 ms given GES in different orientations. The error bars represent the standard errors of the mean. **(d)** and **(i)** show the box plots of the orientation selectivity index (OSI) for neurons selective to each of the four orientations. Grey dots represent individual neurons. **(e)** and **(j)** show the orientation tuning for neurons selective to each of the four orientations. The radii of the concentric circles indicate different firing rates: 0.5, 1, and 1.5 spikes per 100 ms. The black lines connect the firing rates in response to GES with different orientations. Results shown in **(a)** through **(e)** correspond to GES at Target 1. Those in **(f)** through **(j)** correspond to GES at Target 2.

### Orientation selectivity at different post-stimulus latencies

The example in figure 1(e) shows that GES evoked stronger and more reliable neuronal responses at some post-stimulus latencies than others. We asked whether the orientation selectivity was dependent on the response latency. To answer this question, we calculated the PSTH to resolve SUA in time and evaluated the time dependence of orientation selectivity (figure 5). When Target 1 was stimulated in 0°, the time-resolved firing rate (PSTH) of the activated gastric neurons was on average higher around 10 and 43 ms than around 20 ms or after 73 ms (figure 5(b)). The neuronal responses to GES in other orientations did not demonstrate similarly strong time dependence (figure 5(b)).

**Figure 5.**
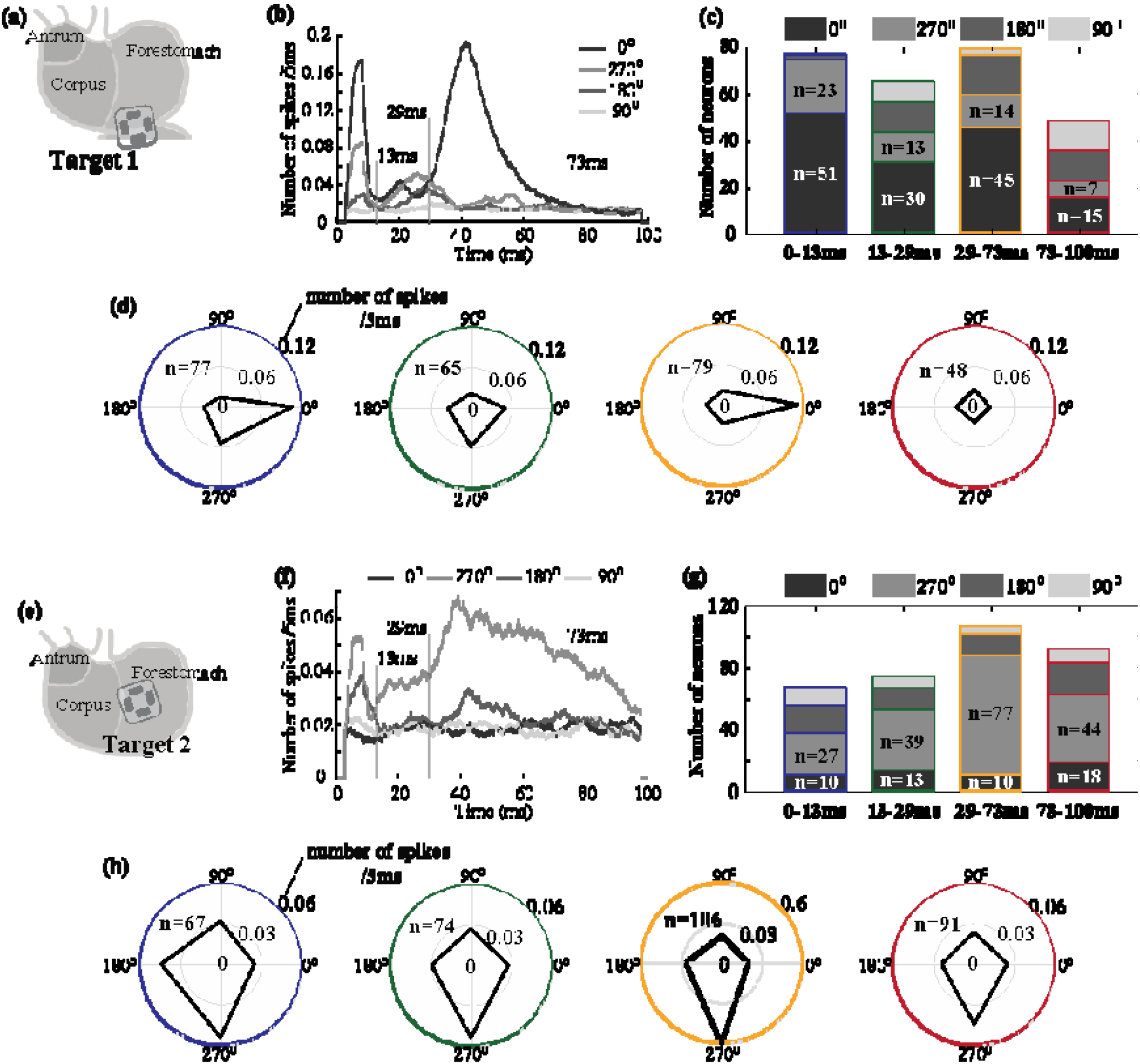
Time dependence of orientation selectivity. For 1-mA GES at Target 1 **(a)** and Target 2 **(e)**, (a) and **(f)** plot the single-unit post-stimulus time histogram (PSTH) averaged across neurons with stimulation applied in 0°, 270°, 180°, and 90°. **(c)** and **(g)** show the number of neurons that preferred each of the four orientations in terms of their responses at four different post-stimulus periods. **(d)** and **(h)** show the number of spikes (coded as the radius) observed during each of the four periods (color-coded as the circles) given GES in each of the four orientations (coded as the angle). Regardless of the stimulus orientation, the total number of neurons activated during each period is denoted as n.

According to the PSTH, we further divided the post-stimulus period into four periods (0-13, 13-29, 29-73, 73-100 ms) and evaluated the orientation selectivity of individual neurons separately for each period. Out of 106 neurons activated by GES at Target 1, 77 neurons were activated from 0 to 13 ms, 65 neurons from 13 to 29 ms, 79 from 29 to 73 ms, and 48 neurons from 72 to 100 ms. For two periods (0-13 ms and 29-73 ms), not only were more neurons activated, but most of the activated neurons were also selective to the same orientation (figure 5(c)). The firing rate of each activated neuron manifested a stronger preference for GES in 0° during the periods of 0-13 ms and 29-73 ms, but much less so for other periods (figure 5(d)). These results suggest that gastric neurons in NTS were selective to the orientation of GES at Target 1 with respect to their responses at two specific delays (about 8 ms and 43 ms) relative to the stimulus onset. For GES at Target 2, the firing rate averaged across all activated neurons was higher when the stimulus was in 270° at nearly all post-stimulus latencies (figure 5(f)). The peak responses were noticeable in two periods (0-13 ms and 29-73 ms). In particular, the period from 29 to 73 ms showed that 77 out of 119 neurons were selective to the same orientation (270°), whereas slightly lower but comparable orientation selectivity was also noticeable for other periods. These results suggest that neurons activated by GES at Target 2 showed consistent selectivity to 270° for all post-stimulus periods, showing a lesser degree of time dependence compared to the responses to GES at Target 1. In summary, when Target 2 is stimulated, more NTS neurons prefer the orientation of 270° within 29-73ms after the stimulus.

### Compare neural responses after the stimulus on different targets

We further asked whether stimulating different gastric locations could activate the same neurons within NTS, and if yes, whether their responses were distinctive and dependent on the stimulus location. To answer these questions, we implanted electrodes to both Target 1 and Target 2 in four rats (figure 6(a)). When applying 1-mA current stimuli in the preferred orientation (0° for Target 1 or 270° for Target 2), we identified the single units that responded to either or both of the locations.

**Figure 6.**
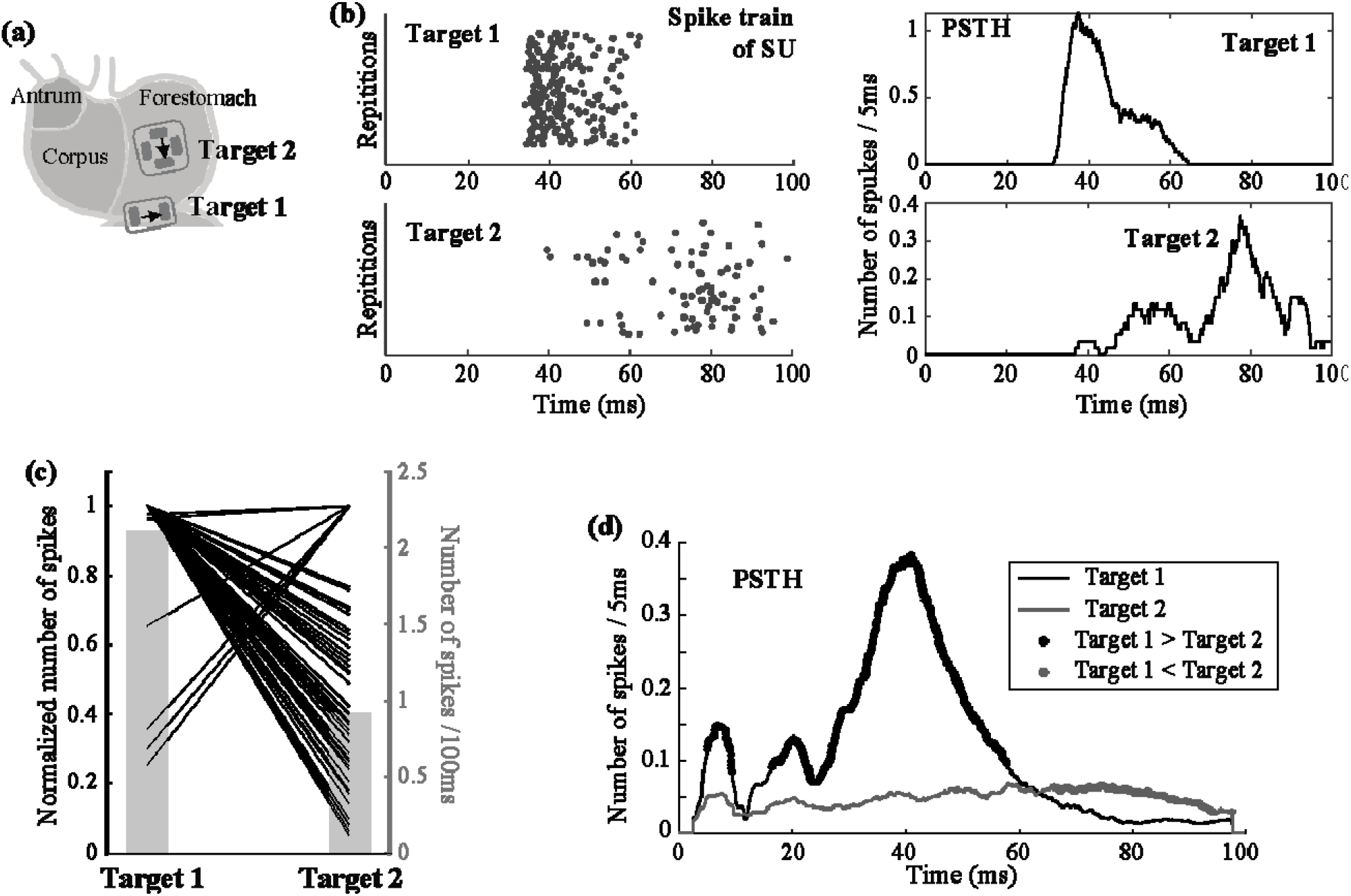
Location dependence of single-unit responses. **(a)** shows 1-mA GES applied in the preferred orientations for both of the two target locations. In **(b)**, the left shows the raster displays of the same neuron’s firing pattern in response to repeated trials of GES at Target 1 (top) and Target 2 (bottom); the right shows the corresponding PSTH. **(c)** plots the relative changes in the time-averaged single-unit responses given GES at Target 1 vs. Target 2. Each line corresponds to a single neuron. For each neuron, the time-average firing rate averaged across trials is always normalized to 1, regardless of whether it is greater when stimulation is at Target 1 or Target 2. The corresponding y-axis is on the left. The grey bar plot shows the averaged firing rate, with the y-axis on the right. **(d)** shows the PSTH averaged across neurons given GES at either Target 1 (black) or Target 2 (gray). The black (or gray) dots highlight the time points where the response to GES at Target 1 (or Target 2) is significantly higher than the response to GES at Target 2 (or Target 1) based on paired t-tests separately evaluated for each time point with □ = 0.05.

In total, 100 neurons were identified: 88 with Target 1, 63 with Target 2, and 51 with both. On average, the single-unit firing rate per 100 ms was 1.72±0.15 (mean ± standard error) for Target 1 and 0.91±0.10 for Target 2. For the 51 neurons responsive to both targets, their average firing rate was 2.12±0.21 for Target 1 and 0.92±0.10 for Target 2. The time-averaged neuronal responses were significantly higher for Target 1 than for Target 2 (paired t-test: t=7.35, p=1.7e-9, dof=50). Only seven neurons showed higher responses to Target 2, whereas the majority showed higher responses to Target 1 (figure 6(c)).

We further resolved the single-unit firing rate in time and averaged the response across the 51 neurons that were responsive to both targets. figure 6(b) shows the firing of a single neuron across repeated stimuli. In this example, the neuron fired at different times in response to the stimulus at Target 1 vs. Target 2. Activation with Target 1 was mostly between 30 and 60 ms, whereas activation with Target 2 was later, especially around 80 ms (figure 6(b)). We averaged the firing rate across neurons, and found that the average neuronal response was higher for Target 1 for the first 60 ms, whereas it was higher for Target 2 between 65 and 100 ms (figure 6(d)).

These results suggest that gastric electrical stimulation selectively applied to longitudinal or circular IMAs at different locations may activate a common group of neurons in NTS but at different times. These neurons are more responsive to stimulation applied to the longitudinal IMAs on the greater curvature at early times (before 60 ms) but are more responsive to the circular IMAs at later times (after 65 ms).

## Discussions

In summary, this paper presents neurophysiological evidence that gastric neurons in NTS are highly selective to the orientation of gastric electrical stimulation applied to the hotspots of longitudinal and circular IMAs in the distal forestomach. To activate gastric neurons in NTS, it is preferable to stimulate the longitudinal IMAs by delivering current pulses to the greater curvature in the longitudinal direction towards the proximal stomach, or to stimulate the circular IMAs by delivering current pulses in between the lesser and greater curvatures in the circular direction towards the greater curvature. This orientation preference is largely generalizable across different stimulus amplitudes (from 0.6 to 1 mA) and dependent on the post-stimulus latency. Further, stimulating longitudinal and circular IMA may activate a shared subset of gastric neurons in NTS, although the activated neurons are relatively more responsive to stimulation of longitudinal IMAs. These findings suggest that gastric neurons in NTS, at least in part, encode orientation-specific muscle activity relayed through vagal afferent neurons that innervate gastric muscle layers through topologically distinctive and orientation selective intramuscular arrays. The scientific and clinical implications of our findings are discussed as below.

The brain sees the gut as a sensory organ (Furness et al., 2013). The primary function of the stomach lies in its motor events effectuated by smooth muscle cells in the gastric wall. The number of gastric smooth muscle cells is orders of magnitude greater than the number of sensory neurons that relay muscle activity to the brain (Powley et al., 2019). The mismatch prevents sensory neurons from uniformly innervating the stomach in a one-to-one manner and necessitates “sparse coding” for efficient gastric representations in NTS. Our results provide functional evidence that NTS encodes and integrates orientation specific muscle activity from the stomach.

Because the stomach moves primarily along two orientations (longitudinal and circular), it makes intuitive sense for NTS to mainly represent longitudinal and circular movements while neglecting other orientations that are less functionally significant (Cao et al., 2019). Such orientation selectivity is enabled by highly specialized morphologies of the afferent nerve terminals (i.e., IMAs) that selectively target longitudinal and circular muscles (Wang and Powley, 2000). As they resemble elongated and parallel arrays, IMAs are uniquely suited to serve as the mechanoreceptors for orientation selective neural coding (Powley and Phillips, 2002).

In addition, the non-uniform distributions of circular and longitudinal IMAs (Powley et al., 2016) may further support sparse coding. Muscle activity is coordinated across regions on the stomach (Lu et al., 2017). Muscle cells are actively coupled to and paced by interstitial cells of Cajal (ICC), and intrinsically modulated by enteric neurons (Furness et al., 2020). Therefore, spatially uniform sampling of muscle activity would otherwise be redundant, rather than sparse. The fact that the IMAs show regional “hotspots” allows the brain to sample only a few regions while likely being able to infer about other regions by reasonable extrapolation given the stomach’s intrinsic organization.

In general, electrical current stimulation is not cell type specific and thus possible to activate other cells, axons, or receptors other than IMAs. Although we did not fully rule out such possibilities, the activation of IMAs is the primary source that drives the orientation selective neural responses at NTS. The current pulses used in this study were too brief and weak to activate smooth muscles. Prior studies suggest that a current pulse should be at least 350 ms and 2 to 4 mA in order to directly cause muscle activation (Du et al., 2009; Hocking et al., 1992; Li & Chen, 2010; McCallum et al., 1998; Tomita, 1966), whereas we used a current pulse of 0.3 ms and 0.6 to 1 mA. In addition, the longitudinal and circular smooth muscles are present in both Target 1 and 2, unlike the differentiated distributions of longitudinal and circular IMAs. If the stimuli were to directly activate muscles, they would have activated both longitudinal and circular muscles and thus could not explain the differential orientation selectivity for GES applied to the two targets.

Arguably, the current pulses might have directly activated the axons of vagal afferents. However, the axonal activation is unlikely to explain the observed orientation preference of afferent responses. Note that axons of vagal afferents run in the circular direction, towards the esophagus and ascending to the brain (Wang & Powley, 2000). If our stimulation were directly applied to the axons, we speculate that the preferred direction would be explainable with the cathode closer to the lesser curvature and the anode closer to the greater curvature, which is opposite to our finding. Our speculation is given the “anodal block” (Brindley & Craggs, 1980; Rijkhoff et al., 1997), which would tend to block action potentials from propagating toward the anode, when peripheral nerve stimulation with bipolar electrodes is delivered from the anode to the cathode. Anodal block has been observed for peripheral nerve stimulation, including the vagus nerve stimulation (Jones et al., 1995; Ahmed et al., 2020; Lu et al., 2020).

Lastly, the current stimuli could also have activated other afferent receptors, e.g., intra-ganglionic laminar endings (IGLE). The morphology of IGLE is like a plate of lamelliform terminal puncta in contact with the ganglion between longitudinal and circular muscle layers (Powley et al., 2019). Such a structure is unlikely to transmit orientation specific signals. Also note that the preferred orientation varies between the two target locations, in agreement with the different regional distributions of IMAs (Powley et al., 2016), but not the distribution of IGLE (Tan et al., 2021).

Results from this study also suggest that neurons in NTS integrate sensory information from different gastric regions. A single neuron could be activated with stimulation applied to different targets and in different orientations. Such a neuron thus has a broad receptive field and a multitude of orientation tuning. Speculatively, the integrative function may arise from second-order neurons that receive converging projections of vagal afferent neurons, as well as higher-order neurons in NTS (Powley, 2000; Whitehead, 1988). The dynamic interplay among different subnuclei may likely account for single-unit responses at different latencies (figure 6). Our speculation agrees with the current understanding that individual subnuclei in NTS do not form a fine-grained topographic map of the stomach, but progressively integrates sensory input (Powley, 2000; Rogers et al., 1995; Travagli & Anselmi, 2016).

The orientation selectivity of neurons in NTS is likely to have a profound implication to where and how GES should be applied. To the best of our knowledge, prior studies on characterizing the effects of GES with various parameters have rarely addressed the effects of the stimulus orientation, except for one preliminary study (Cao et al., 2019). In clinic practice, GES is applied to the corpus or antrum (Abell et al., 2003; Hasler, 2009). This choice was perhaps historical, dating back to the initial attempt of using direct currents to pace gastric contractions (Kelly & La Force, 1972). While gastric pacing has not been widely adopted (Hasler, 2009; Yin, & Chen, 2008), the primary goal of GES has focused on modulating neural circuits that control the stomach. However, clinical applications with GES largely or entirely ignore the morphological, topological or chemical phenotypes of different kinds of nerve terminals. The translational potential of our findings is obscured by the anatomical difference between rats and humans. The extrinsic innervation has been extensively mapped out for the rat stomach (Powley et al., 2019), but remains under-studied in humans. It remains unclear whether and to what extent our findings in rats are generalizable to humans. However, humans also have mechanical afferent receptors in the proximal stomach that are sensitive to changes in gastric volume (Brookes et al., 2013; Carmagnola et al., 2005; Penagini et al., 2004). It is speculatively plausible that gastric neurons in human NTS may also be selective to the orientation of gastric electrical stimulation applied to intramuscular nerve endings of afferent neurons. However, specific locations for orientation-specific stimulation remain unclear and await comparative studies of the human vs. rat anatomy.

The present study limits its scope to neurons in NTS. However, it is likely that the effects of orientation selective afferent stimulation may extend beyond NTS and give rise to downstream changes in both the gut and the brain. Neurons in NTS project to the dorsal motor nuclei of the vagus, from which vagal efferent nerves innervate the gut (Travagli & Anselmi, 2016). It is reasonable to speculate that orientation and location-specific GES may selectively activate the vagovagal reflexes to regulate gastric physiology, such as gastric secretion and motility (Travagli et al., 2006; Browning & Carson, 2021). Further, gastric neurons in NTS ascend projections to the forebrain (Browning & Travagli, 2011), likely affecting gastric interoception and eating behaviors (Babic & Browning, 2014; Chen, 2004; Han et al., 2018). Future studies are required to address these potential peripheral and central effects in order to further understand the functional significance of orientation-selective gastric electrical stimulation.

In conclusion, results from this study suggest that the orientation is another key parameter for GES, in addition to other parameters, e.g., pulse width, amplitude, or frequency. The orientation of stimulation is worth considering in clinical practice and further exploration in future studies. For GES to activate gastric neurons in NTS, two effective locations are in the distal forestomach: one on the greater curvature and the other in between the lesser and greater curvatures, as informed by the regional distributions of IMAs. For the former, the optimal orientation is the longitudinal direction towards the distal stomach. For the latter, the optimal orientation is the circular direction towards the greater curvature. GES applied with such optimal locations and orientations may trigger a cascade of integrative processing within and beyond NTS, and likely give rise to therapeutic benefits. This expected outcome remains speculative and awaits confirmation in future preclinical or clinical studies.

## Acknowledgment

The authors thank Dr. Robert Phillips for his guidance on the electrode implementation for gastric electrical stimulation, thank Drs. Zhenjun Tan, Kun-Han Lu for initial discussions in this project, and thank Drs. Mark Sayles and Xueguo Zhang for their initial assistance in the electrophysiological recording. This study is supported by NIH grants: OD023847 (TP), OD030538 (ZL), AT011665 (ZL).

## Ethical statements

This study was approved by the Institutional Animal Care and Use Committee and the Laboratory Animal Program at Purdue University.

